# A hippocampal-hypothalamic circuit essential for anxiety-related behavioral avoidance

**DOI:** 10.1101/2022.01.17.476545

**Authors:** Jing-Jing Yan, Ai-Xiao Chen, Wen Zhang, Ting He, Xiao-Jing Ding, Zi-Xian Yu, Yan-Li Zhang, Mengge He, Haohong Li, Xiao-Hong Xu

**Affiliations:** Institute of Neuroscience, State Key Laboratory of Neuroscience, CAS Center for Excellence in Brain Science and Intelligence Technology, Chinese Academy of Sciences, Shanghai 200031, China; Shanghai Center for Brain Science and Brain-Inspired Intelligence Technology, Shanghai 200031, China; University of Chinese Academy of Sciences, Beijing 100049, China; The MOE Frontier Research Center of Brain & Brain machine Integration, Zhejiang University School of Brain Science and Brain Medicine, Hangzhou Zhejiang 310058, China; Britton Chance Center for Biomedical Photonics, Wuhan National Laboratory for Optoelectronics, Huazhong University of Science and Technology, Wuhan, Hubei 430074, China; MoE Key Laboratory for Biomedical Photonics, Collaborative Innovation Center for Biomedical Engineering, School of Engineering Sciences, Huazhong University of Science and Technology, Wuhan, Hubei 430074, China

## Abstract

Anxiety over perceived threats triggers avoidance behavior, but the underlying neural circuit mechanism remains poorly understood. Taking hints from the deep connection between anxiety and predator defense, we examined the role of the anterior hypothalamic nucleus (AHN), a critical node in the predator defense network, in anxiety-related behaviors. By recording Ca^2+^ transients in behaving mice, we found that activity of AHN GABAergic (AHN^Vgat+^) neurons showed individually stable increases when animals approached unfamiliar objects in an open field (OF) or explored the open arm of an elevated plus-maze (EPM). Moreover, AHN^Vgat+^ neuron activity foreshadowed behavioral retreats and correlated with object and open-arm avoidance. Crucially, exploration-triggered optogenetic inhibition of AHN^Vgat+^ neurons dramatically reduced avoidance behaviors. Furthermore, retrograde viral tracing identified the ventral subiculum (vSub) of the hippocampal formation as a significant input to AHN^Vgat+^ neurons in driving avoidance behaviors. Thus, the activity of the hippocampal-hypothalamic pathway promotes idiosyncratic anxiety-related behavioral avoidance.

## Introduction

Anxiety represents an emotional state of apprehension about remote, potential, unpredictable, or ill-defined threats ^1–4^. It keeps individuals vigilant about potential harms, thereby preparing them for safety measures ^4, 5^. Using behavioral tests that exploit the “approach-avoidance” conflict ^6^, such as the open field test and the elevated plus-maze (EPM), previous studies have identified many brain areas that work in concert to regulate approach-avoidance behaviors ^7–10^. Some of these brain regions, such as ventral CA1 (vCA1) of the hippocampus, the lateral septum nuclei (LS), and the bed nucleus of the stria terminalis (BNST), modulate approach-avoidance behaviors in part through projections to hypothalamic nuclei ^11–13^. Intriguingly, brief predator encounters increase anxiety levels in species ranging from flatworms to fish, rodents, and primates ^14–17^, pointing to an evolutionarily conserved mechanism linking predator-provoked defensive behavior with anxiety ^18–21^. Conversely, animals selectively bred for high anxiety traits show increased defensive avoidance to predator cues ^22^. Moreover, anti-anxiety drug treatments diminish predator defense in normal animals ^23, 24^. Together, these findings suggest that neural substrates underlying anxiety-related behaviors overlap with those mediating predator defense behavior.

The present study focused on the anterior hypothalamic nucleus (AHN), which reciprocally connects with the ventromedial hypothalamus (VMH) and the dorsal premammillary nucleus of the hypothalamus (PMd) to form the hypothalamus predator defense network ^25, 26^. Predator cues activate this network, particularly VMH ^26–30^. Optogenetic activation of VMH neurons or their projections to AHN is sufficient to drive avoidance behaviors such as flight ^31^. However, lesioning the AHN failed to produce an effect on predator defense as clear as that of VMH or PMd ^27, 28, 32, 33^. By comparison, anti-anxiety drug treatment reduces predator-induced c-Fos signals in AHN but not VMH ^24^. Additionally, LS neurons that express type 2 corticotropin-releasing factor receptor (Crfr2) enhance stress-induced anxiety behaviors and cortisol release through projections to AHN ^12^. Based on these results, we focused on AHN neurons as a potential convergence site for neural circuits linking anxiety with threat-evoked avoidance behaviors.

We first found that activity of AHN GABAergic neurons (AHN^Vgat+^) strongly correlated with the mouse avoidance behaviors in two standard anxiety tests, with each mouse exhibiting consistent and individual-specific AHN^Vgat+^ activity changes. Furthermore, we showed that optogenetic inhibition of AHN^Vgat+^ neurons at the time of exploration reduced subsequent avoidance behaviors. Using pseudorabies virus retrograde tracing, we further identified the ventral subiculum (vSub) of the hippocampal formation as a major input to AHN^Vgat+^ neurons in driving avoidance behaviors. These results point to the importance of the hippocampal-hypothalamic circuit in controlling anxiety-related behavioral avoidance.

## Results

### Strong temporal correlation of AHN^Vgat+^ neuron activity with anxiety-related avoidance behavior in a modified open field paradigm

Center avoidance and peripheral preference in an open field test are behavioral parameters that indicate rodent anxiety levels ^6^. By introducing an unfamiliar object (a battery) to the center of an open field ∼10 mins after a mouse freely explored the arena, we found that this procedure led to more substantial center avoidance and peripheral preference (**Fig. 1a**), indicating that an unfamiliar object elevates the anxiety level. Such behavioral changes were not observed in control animals that were allowed to explore the open field continuously for 20 mins, with the experimenter’s hand interruption briefly without placing the object (**Fig. 1b**). Thus, object-evoked behavioral changes were unlikely caused by fatigue, habituation, or human interference.

**Fig. 1.**
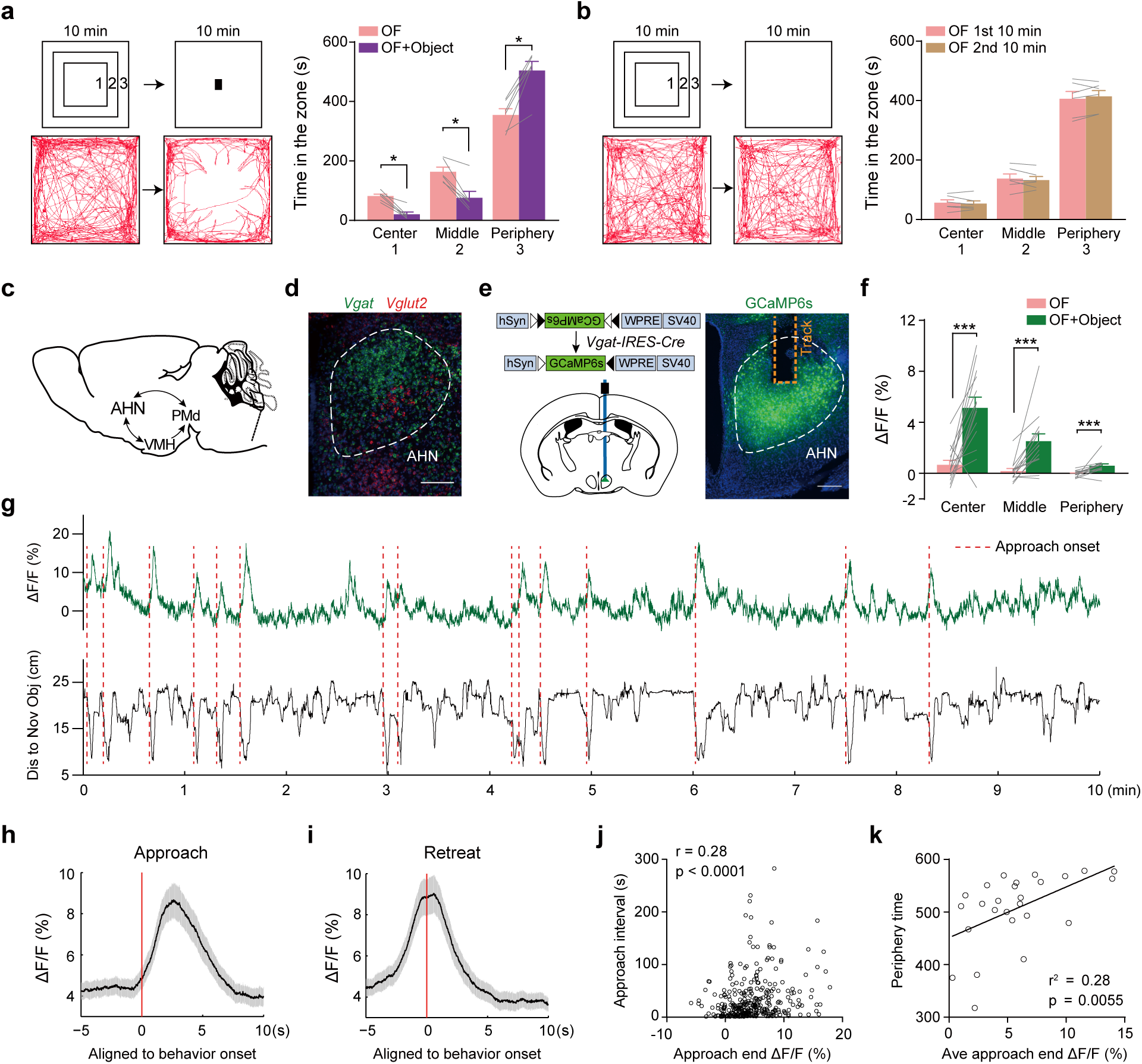
Strong temporal correlation of AHN^Vgat+^ neuron activity with anxiety-related avoidance behavior in a modified open field paradigm. (**a**) The open field test modified with the introduction of an unfamiliar object 10 mins after the initial exploration. “1”, “2” and “3” denote the “center”, “middle” and “peripheral” zone of the open field. The example trajectory (bottom) and the quantification (right) show that animals spent more time in the peripheral zone away from the center after object introduction. n = 6 mice. (**b**) Control assays in which animals were allowed to explore the open field continuously for 20 mins. The example trajectory and the quantification (right) show similar time spent in all three zones in the first and second 10 mins of the open field test. n = 6 mice. (**c**) Schematic illustration of the “hypothalamus predator defense circuit”. (**d**) A representative image showing the fluorescent *in situ* signals of *Vgat* and *Vglut2* mRNA in AHN. Scale bar, 200 μm. (**e**) Left, the strategy to monitor GCaMP6s signals in AHN^Vgat+^ neurons. Right, a representative image showing restricted GCaMP6s expression in AHN. Scale bar, 200 μm. (**f**) Average ΔF/F values detected in the “center”, “middle” and “periphery” zone before and after object introduction in GCaMP6s animals. n = 14 mice. (**g**) A representative trace of ΔF/F signals (green, top) aligned to the relative distance (black, bottom) between a GCaMP6s animal and the object. Red dashed lines denote onset of approach bouts. (**h-i**) Average ΔF/F values of GCaMP6s signal aligned to approach (**h**) or retreat onset (**i**) at the time “0”. Shades indicate the SEM. (**j**) Correlation between the GCaMP6s ΔF/F value at the end of an approach and the latency to initiate the following approach. n = 351 bouts from 14 mice. (**k**) Correlation between average approach-end GCaMP6s ΔF/F value and the time animals spent in the periphery zone. n = 26 trials from 14 mice. *, p < 0.05; ***, p < 0.001.

Because behavioral changes elicited by an unfamiliar object in the open field test are similar to those caused by brief predator exposure (Kennedy et al., 2020), we further inquired whether the hypothalamus predator defense circuit, particularly AHN, was engaged during the process (**Fig. 1c**). First, our *in situ* mapping of mRNAs of vesicular transporters for GABA and glutamate (*Vgat* and *Vglut2*) showed that, among all *Vgat+* and *Vglut2+* neurons in AHN, the vast majority (83.0 ± 5.7%, n = 3 mice) expressed *Vgat* (**Fig. 1d)**. We thus used the *Vgat-IRES-Cre* line to target AHN *Vgat+* (AHN^Vgat+^) neurons. We independently validated the fidelity of this mouse line by injecting adeno-associated virus (AAVs) encoding Cre-inducible EYFP into AHN, in which we found 98.9 ± 0.7 % of GFP+ neurons expressed *Vgat* (Extended Data Fig. 1a, n = 3 mice).

To monitor the activity of AHN^Vgat+^ neurons, we injected AAVs encoding Cre-inducible GCaMP6s, or EYFP as the control, into AHN of *Vgat-IRES-Cre* mice and implanted an optic fiber above the injection site (**Fig. 1e**). These procedures did not result in apparent changes in object-evoked avoidance behavior in the open field (before vs. after object introduction, peripheral zone time, 385.8 ± 10.8 s vs. 500.7 ± 16.3 s, p < 1×10^-4^, n = 22 mice). Before object introduction, GCaMP6s signals of AHN^Vgat+^ neurons were not significantly modulated by the location of the animal in the open field (**Fig. 1f**). Remarkably, after object introduction, GCaMP6s signals of AHN^Vgat+^ neurons elevated considerably, with the most dramatic increase observed when the mouse arrived at the open field center zone (**Fig. 1f**). Notably, such fluorescent signal changes were not observed in control animals that expressed EYFP in AHN^Vgat+^ neurons (Extended Data Fig. 1b-d), indicating that changes in GCaMP6s signals were unlikely caused by motion artifacts.

Moreover, we found that AHN^Vgat+^ GCaMP6s signals tracked with the animal’s distance relative to the object (**Fig. 1g**), ramping up as the animal approached the object and down as it retreated to the peripheral zone (**Fig. 1h-i**). For individual trials, the overall temporal dynamics of AHN^Vgat+^ fluorescence signals strongly correlated with “approach-retreat” bouts in GCaMP6s mice with an average correlation coefficient (r) of 0.28 ± 0.05 (n = 14 mice), significantly higher than that of EYFP control mice (r = -0.03 ± 0.01, n = 8 mice; p < 1×10^-4^). Furthermore, the peak value of AHN^Vgat+^ GCaMP6s signals at the end of a center approach, a turning point before the retreat, positively correlated with the latency to initiate the next approach (**Fig. 1j**, r = 0.28, p < 1×10^-4^), suggesting that close encounter with the object elevated the anxiety. Along the same line, the average “turning point” AHN^Vgat+^ GCaMP6s signals in a trial significantly correlated with the total duration that the animal spent in the peripheral zone away from the object (**Fig. 1k**, r^2^ = 0.28, p = 0.0055).

As a comparison study, we placed an object in the mouse’s home cage for three days for familiarization and then performed the open field test with the familiarized object in the center (Extended Data Fig. 2a). Interestingly, despite intense object investigation, AHN^Vgat+^ GCaMP6s signals did not change during approach or retreat (Extended Data Fig. 2b-d). No signal was observed when the mouse investigated, sniffed, or mounted a female mouse introduced to its homecage either (Extended Data Fig. 2e-h). Thus, AHN^Vgat+^ neuron activity does not reflect exploratory actions or social activity. Together, these results show a robust temporal correlation between AHN^Vgat+^ neuron activity and anxiety-related avoidance behavior.

### Object-evoked AHN activity shows individual specificity and converges with predator cue response

To investigate whether any specific features of the object used (a battery) is responsible for evoking AHN^Vgat+^ activity, we performed a new set of experiments using three other alternative items, an acrylic cuboid cube, a toy airplane, and a metal paper clip, in addition to the battery, as the unfamiliar object (Extended Data Fig. 3a). We individually presented these four objects on separate testing days in a pseudo-randomized order (Extended Data Fig. 3b). We found that all objects drove the tested animals to spend more time in the peripheral zone after being introduced to the open field (Extended Data Fig. 3c). Furthermore, we found a similar temporal correlation of ramping AHN^Vgat+^ GCaMP6s signals with approach-retreat bout and with the time spent in the peripheral zone for all four objects (**Fig. 2a-b**). Notably, the AHN^Vgat+^ GCaMP6s signals and avoidance behavior evoked by the unfamiliar object were variable among different mice, yet the same mouse showed highly consistent responses towards different objects. Further pair-wise analysis showed a strong correlation of data between individual trials of two different objects, for the average turning point GCaMP6s signals and the time spent in the peripheral zone (**Fig. 2c**). This individual specificity further supports the notion that elevated AHN^Vgat+^ neuron activity underlies anxiety-related avoidance behavior.

**Fig. 2.**
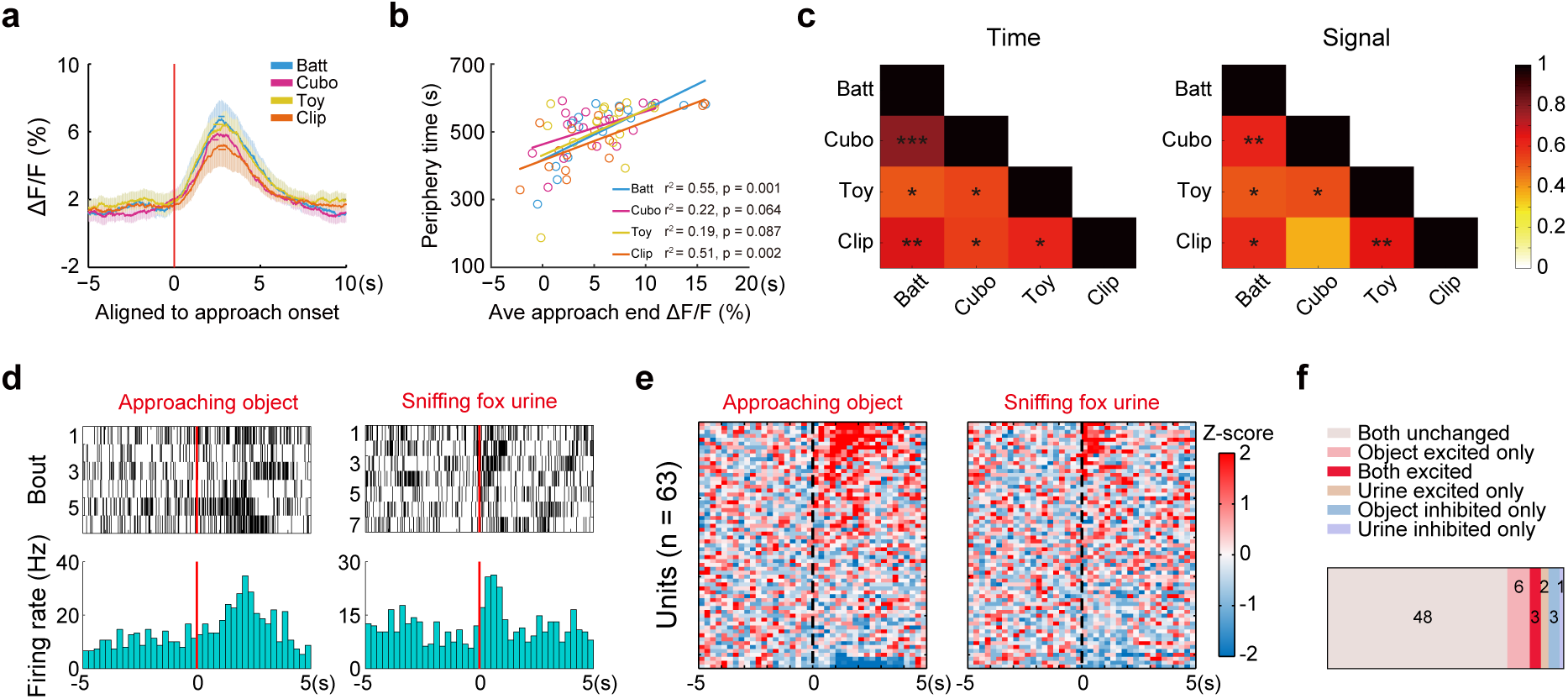
Object-evoked AHN activity shows individual specificity and converges with predator cue response. (**a**) Average ΔF/F signals ± SEM (shades) aligned to approach onset toward different unfamiliar objects. The colored bars show the average retreat onset ± SEM. (**b**) Correlations between average approach-end ΔF/F value and the time spent in the open field periphery zone after the introduction of different unfamiliar objects. n = 16 mice. (**c**) Pair-wise correlations between the time spent in the periphery zone (left) and the average approach-end ΔF/F value (right) across the four object conditions. The heat map (scale on the right) represents the correlation co-efficiency (r) value with the p values, indicated by stars, depicted in each cell for each pair. (**d-f**) Single unit recordings of AHN neurons. (d) Raster plot (top) and the average (bottom) of the firing of an example single-unit aligned to the onset of object-approaching behavior (left) and fox urine sniff (right). (e) Heatmap representation of the normalized single-unit responses in Z scores sorted by response magnitude aligned to behavioral onset. n = 63 units from 3 mice. (f) Quantification of the number of single units in each response category. *, p < 0.05; **, p < 0.01; ***, p < 0.001.

To examine whether object-activated AHN neurons converge with those responding to predator cues, we performed single-unit recordings in AHN while sequentially exposing the mouse to an unfamiliar object in an open field and a piece of paper spotted with fox urine in a clean cage (Extended Data Fig. 4, **Fig. 2d-f**). 9 out of the 63 single units recorded from three mice increased firing during object approach, and 5 increased during fox urine sniff (**Fig. 2e)**. Moreover, 3 of these units responded to both object and fox urine (**Fig. 2d, e)**. Thus, object and predator cues activated partially overlapping AHN neuronal ensemble. These results further support elevated AHN neuron activity as a neural mechanism linking anxiety with hardwired avoidance behaviors evolutionarily selected for predator defense.

### Inhibiting object-evoked AHN^Vgat+^ activity reduces avoidance

We next examined whether inhibiting AHN^Vgat+^ neuron activity evoked by an unfamiliar object could abolish object-induced increases in anxiety and avoidance behavior. To this end, we bilaterally injected AAVs encoding Cre-inducible GtACR1, or EYFP as the control, into AHN of *Vgat-IRES-Cre* male mice (**Fig. 3a**) and implanted an optic fiber 300-500 μm above each injection site (**Fig. 3b**). We used *ex vivo* patch-clamp recordings to confirm that pulses of blue light (473nm, 20ms, 20Hz) effectively and reversibly silenced GtACR1-expressing AHN^Vgat+^ neurons (**Fig. 3c-d**). By analyzing fiber-photometry recorded animals (**Fig. 1**), we found that the starting point for approach bouts toward the object was mostly located within the peripheral zone (**Fig. 3e**). Therefore, we delivered light pulses whenever the mouse left the peripheral zone after object introduction to inhibit object-evoked AHN^Vgat+^ neuron activity during the approach (**Fig. 3e**). These light pulses had no effect in control EYFP mice but completely abolished the object avoidance and peripheral preference in GtACR1 mice (**Fig. 3f-g**). Thus, the elevated activity of AHN^Vgat+^ neurons during the approach could reflect the increased anxiety level caused by the object.

**Fig. 3.**
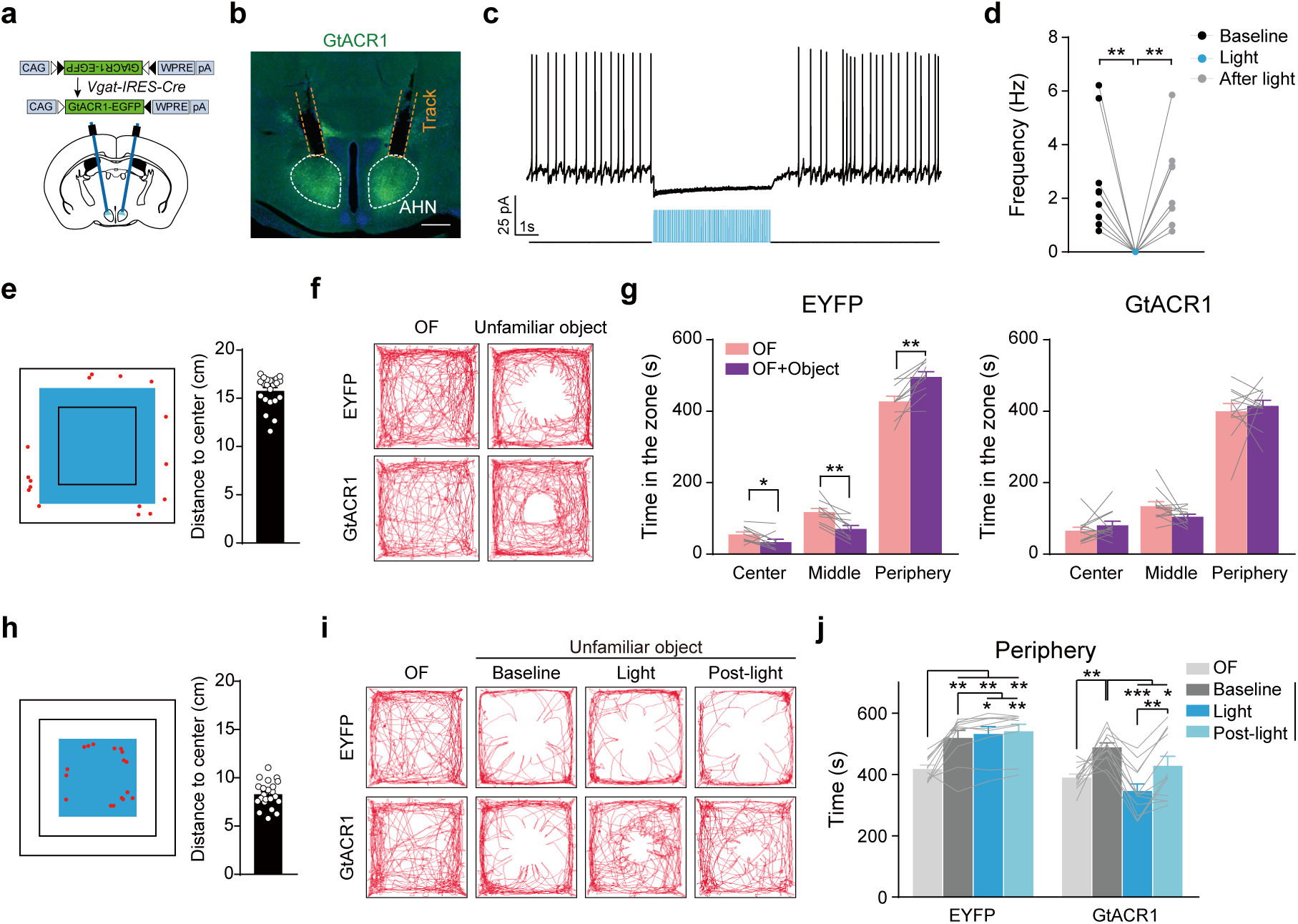
Optogenetic inhibition of object-evoked AHN^Vgat+^ activity reduces avoidance. (**a**) The viral strategy to optogenetically inhibit AHN^Vgat+^ neurons. (**b**) A representative *post-hoc* image showing GtACR1 expression in AHN and tracks of fibers implanted above. Scale bar, 200 μm. (**c-d**) A representative trace (c) and quantifications (d) show light-mediated inhibition of GtACR1-expressing neurons. **(e-j)** Optogenetic inhibition of AHN^Vgat+^ neurons. (e) & (h) Light delivery pattern shown by the blue square for experiments in (f-g) & (i-j) respectively. Red dots denote the starting location of all approach (e) or retreat (h) bouts in a representative trial. The quantification on the right shows the average distance (in the vertical or horizontal direction) between approach (e) or retreat (h) starting location and the open field center, where the object was placed. n = 22 mice. (f) & (i) Representative trajectories of an EYFP or GtACR1 male with light delivered in the center and middle zone (f) or center zone only (i) after object introduction. (g) & (j) Quantification of the time spent in the indicated zone before or after object introduction in EYFP (n = 10) and GtACR1 males (n = 12). *, p < 0.05; **, p < 0.01; ***, p < 0.001.

Because the activity of AHN^Vgat+^ neurons climaxed before the retreat, we further inhibited these neurons with more precise temporal control by applying the light pulses when mice arrived at the center zone where the unfamiliar object was placed, and most retreat bouts were initiated (**Fig. 3h**). For this set of experiments, we recorded baseline behavior for 10 min before and after the introduction of an object (a battery or a cuboid) to the center (**Fig. 3i**), and the light was then delivered whenever the mouse arrived at the center zone during the next 10 mins (**Fig. 3i**). Remarkably, optogenetic inhibition of AHN^Vgat+^ neurons in the center zone drastically reduced the object avoidance, as shown by more time spent in the center zone and less time in the peripheral zone during the inhibition phase as compared to that during the baseline period (**Fig. 3i-j,** Extended Data Fig. 5a-b). This behavioral effect even persisted after the cessation of the light (**Fig. 3i-j,** Extended Data Fig. 5a-b).

In the above experiments, the extended duration the mice spent in the presence of the object (30 min) by itself did not reduce object avoidance since EYFP control mice showed no reduction of object avoidance before and after light stimulation (**Fig. 3j,** Extended Data Fig. 4a-b). Furthermore, the behavioral effects of optogenetic inhibition in GtACR1 animals were unlikely due to a light-conditioned place preference (CPP). When we paired light delivery to one of the two chambers in a CPP apparatus (Extended Data Fig. 5c), light did not lead to preference of the paired chamber in either EYFP or GtACR1 animals (Extended Data Fig. 5d). Together, these results support that elevated AHN^Vgat+^ neuron activity underlies object-induced anxiety and avoidance behavior.

### Progressive engagement of AHN^Vgat+^ neurons during the elevated plus-maze test

To examine whether AHN^Vgat+^ neurons regulate anxiety-related behaviors in another scenario, we monitored the activity of AHN^Vgat+^ neurons in mice exploring an elevated plus-maze (EPM), where avoidance of the open arm indicates general anxiety levels of the mice. In general, we found that the mouse exhibited significantly higher AHN^Vgat+^ neuron activity in the open arm than the closed arm, as shown by the heat map of recorded activity from an example mouse (**Fig. 4a**). For all mice recorded, the average GCaMP6s signal (ΔF/F) in the open arm (2.7 ± 0.8%) was significantly higher than that found in the closed arm (-0.5 ± 0.1%, n = 14 mice, p = 0.0028). No difference of signals was found in EYFP control mice (open arm 0.4 ± 0.4%, closed arm, -0.1 ± 0.1%, n = 8 mice, p = 0.74).

**Fig. 4.**
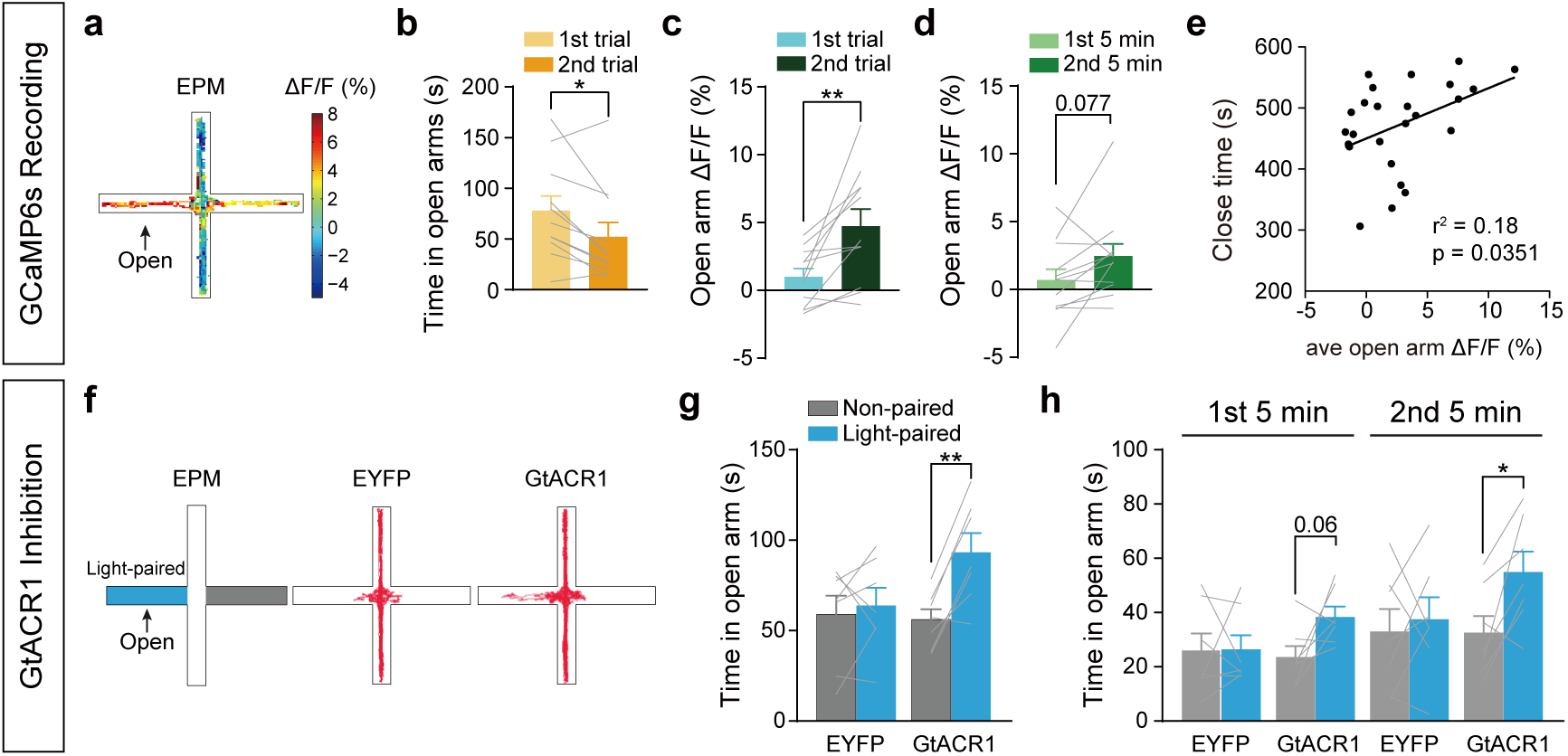
Progressive engagement of AHN^Vgat+^ neurons on EPM. (**a-e**) Recording of AHN^Vgat+^ GCaMP6s signals on EPM. (a) Heatmap representation of EPM ΔF/F value in an example trial. (b-c) Average open-arm time (b) and ΔF/F values (c) in the first or second trial. n = 11 mice. (d) Average ΔF/F values in the first or second 5 min of the first trial. n = 12 mice. (e) Correlation between average open arm ΔF/F value and the time spent in the closed arm. n = 14 mice. (**f-h**) GtACR1-mediated optogenetic inhibition of AHN^Vgat+^ neurons on EPM. (f) Left, schematics showing light delivery restricted to a random open arm; right, example trajectories from an EYFP or a GtACR1 animal as indicated. (g-h) Time spent in open arm. n = 7 EYFP and 7 GtACR1 males. *, p < 0.05; **, p < 0.01.

Mice exhibit progressive higher anxiety during repeated exposure to EPM ^34^. This was supported by our observation of a marked reduction in open arm exploration in the second trial compared to the first trial (**Fig. 4b**). Notably, we found significantly higher AHN^Vgat+^ activity in the open arm during the second trial than during the first trial (**Fig. 4c**). A progressive increase in AHN^Vgat+^ activity could be discerned even within the first trial, as shown by higher activity during the second 5 min than during the first 5 min of the trial (**Fig. 4d**). Correlation analysis showed that for all trials, the average open-arm GCaMP6s signals positively correlated with the total time that the mouse spent in the closed arm (**Fig. 4e**). Furthermore, we found that the AHN^Vgat+^ activity in the open arm of EPM significantly correlated with object-evoked AHN^Vgat+^ activity at the open field center (r^2^ = 0.30, p < 0.044), suggesting that the elevation of AHN^Vgat+^ neuron activity may play a similar role in these two different anxiogenic situations.

To further determine whether the activity of AHN^Vgat+^ neurons is critical for EPM open arm avoidance, we optogenetically inhibited these neurons by virally expressing GtACR1 in AHN^Vgat+^ neurons and applied blue light via implanted optic fibers. The light was applied to only one of the two open arms (**Fig. 4f**). We found that GtACR1 mice spent significantly more time exploring the light-illuminated open arm, while EYFP control mice spent a comparable amount of time in either open arm (**Fig. 4f-g**). Interestingly, this behavioral effect was more substantial during the second 5 min than the first 5 min of the trial, consistent with the progressive increase of AHN^Vgat+^ neuron activity described above (**Fig. 4h**). Furthermore, we found a similar increase in open arm exploration in a new set of experiments when we shined the light to both open arms to inhibit AHN^Vgat+^ neurons (Extended Data Fig. 6). Together, these results indicate that AHN^Vgat+^ neuron activity is essential for EPM open arm avoidance.

### Hippocampal formation sends monosynaptic excitatory inputs to AHN^Vgat+^ neurons

We next sought to identify the synaptic inputs that drive AHN^Vgat+^ neuron activity and avoidance behavior in anxiety-provoking situations, using a pseudorabies virus tracing strategy ^35^. A mixture of AAVs encoding Cre-inducible avian retroviral receptor (TVA)-GFP and rabies glycoprotein (RG) was unilaterally injected into AHN of *Vgat-IRES-Cre* male mice, followed three weeks later with the injection of EnVA-coated pseudorabies virus expressing dsRed but lacking the glycoprotein into the same site (**Fig. 5a**). Our results showed many GFP+/dsRed+ ‘‘starter’’ cells in AHN (**Fig. 5b**) and retrograde-labeled dsRed+ cells in many upstream brain regions (**Fig. 5c**, Extended Data Fig. 7a-b). For parallel controls, mice were injected with AAVs encoding Cre-inducible TVA but not RG (Extended Data Fig. 7c), a procedure preventing the spread of pseudorabies virus after infection of the “starter” cells. We found no dsRed+ cells in upstream brain regions of these control mice (Extended Data Fig. 7c-e), validating the retrograde viral tracing strategy. Quantification of labeled upstream neurons showed that AHN^Vgat+^ neurons received significant inputs from the lateral septum (LS), medial preoptic area (MPO), and bed nucleus of stria terminalis (BNST) (Extended Data Fig. 7b), all of them are known to harbor predominantly GABAergic neurons (www.mouse.brain-map.org). On the other hand, among upstream regions likely to provide excitatory inputs to AHN^Vgat+^ neurons, the ventral subiculum (vSub) of the hippocampal formation, which has been implicated in stress response, emotion regulation, and spatial navigation ^36^, had the highest percentage of retrogradely labeled neurons (**Fig. 5d**).

**Figure 5.**
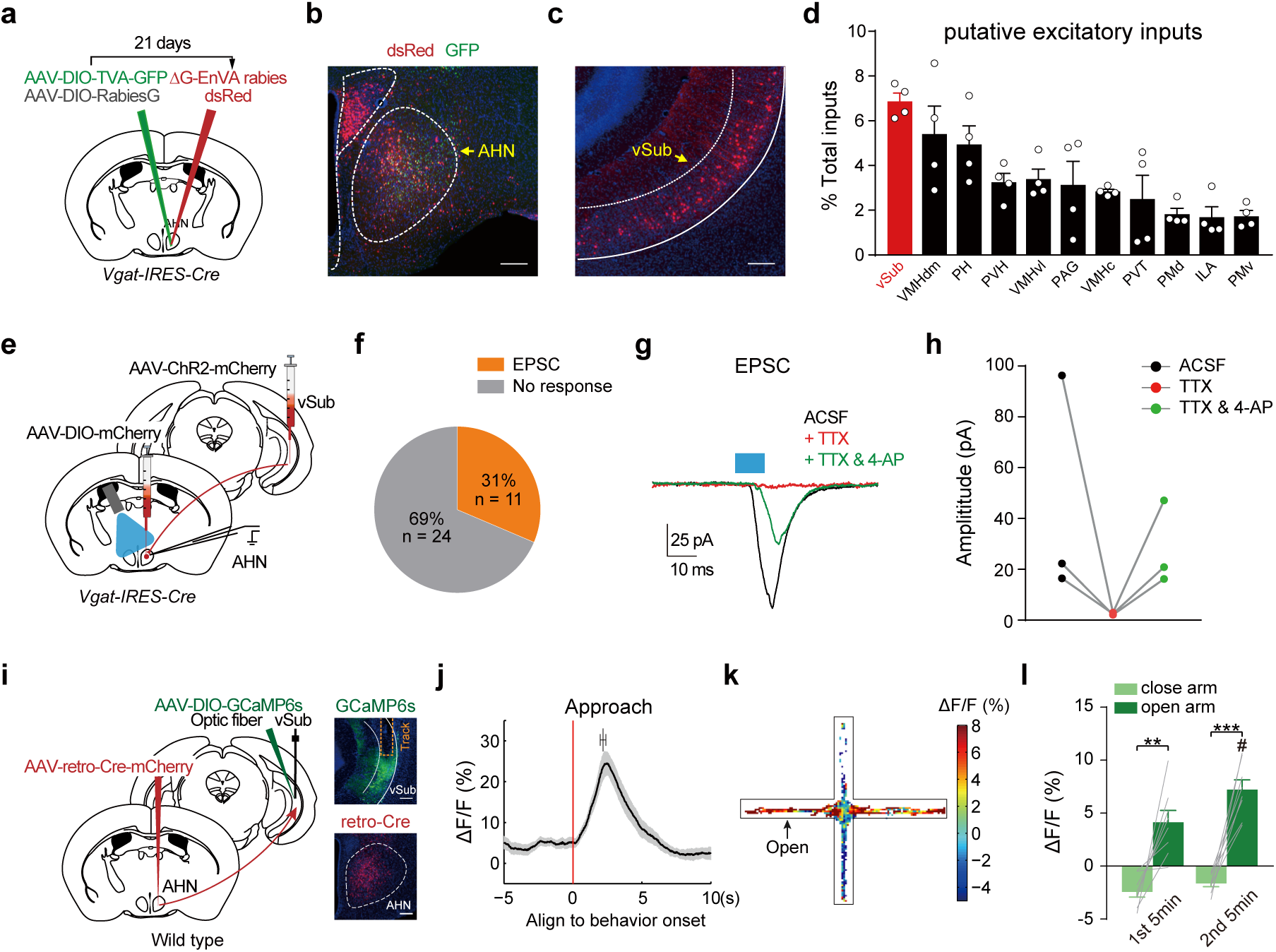
Hippocampal formation sends monosynaptic excitatory inputs to AHN^Vgat+^ neurons. (**a-d**) Retrograde tracing of inputs to AHN^Vgat+^ neurons. (a) Schematics of the viral strategy. (b-c) A representative image showing infection of AHN^Vgat+^ neurons by AAV-DIO-TVA-GFP and EnVA-pseudotyped rabies virus expressing dsRed (b), and retrograde-labeled dsRed+ cells in vSub (c). Scale bar, 200 μm. (d) Quantification of dsRed+ neurons in candidate excitatory brain areas as the percentage of total dsRed+ cells detected outside AHN. n = 4 mice. (**e-h**) Validation of vSub inputs to AHN^Vgat+^ neurons as monosynaptic and excitatory *via* patch clamp. (e) Schematics of the viral and electrophysiological recording strategy to probe vSub inputs to AHN^Vgat+^ neurons. (f) The number and percentage of recorded neurons that showed light-evoked EPSC. n = 35 cells from 6 animals. (g-h) Example traces (g) and quantifications (h) of light-evoked EPSC amplitude in AHN^Vgat+^ neurons under different conditions. Blue bar indicating light pulse stimulation (10 ms). (**i-l**) Recording the activity of AHN-projecting vSub neurons. n = 8. (i) Left, schematics of the viral strategy to target AHN-projecting vSub neurons retrogradely. Right, representative images showing GCaMP6s expressed in vSub and the track of the implanted fiber above (top), and retro-Cre expression in AHN (bottom). Scale bar, 200 μm. (j) Average of GCaMP6s ΔF/F signals ± SEM (shades) in AHN-projecting vSub neurons aligned to approach onset. The bar on top shows average retreat onset ± SEM. (k) Heatmap depiction of EPM GCaMP6s ΔF/F signals in an example trial. (l) The average ΔF/F values in the closed arm and open arm in the first and second 5 min of the trial. “#” denotes significant differences between open arm ΔF/F values in the first and second 5 min (p < 0.01). **, p < 0.01; ***, p < 0.001.

To confirm the monosynaptic connectivity from vSub neurons to AHN^Vgat+^ neurons, we injected AAVs encoding hSyn-ChR2-mCherry unilaterally into vSub and AAVs encoding Cre-inducible mCherry into AHN of *Vgat-IRES-Cre* mice to label AHN^Vgat+^ neurons fluorescently. We then performed patch-clamp recording from AHN^Vgat+^ neurons in acute brain slices containing AHN to monitor synaptic activity evoked by vSub projections (**Fig. 5e**). Of 35 mCherry-expressing AHN^Vgat+^ cells recorded from 6 mice, single light pulses (473 nm, 10 ms) evoked excitatory postsynaptic currents (EPSCs) in 11 cells (**Fig. 5f & g**), with a connection rate of 31%. The amplitude and latency for light-evoked EPSCs were 21.9 ± 7.8 pA and 5.5 ± 0.3 ms, respectively. Furthermore, tetrodotoxin (TTX) blocked light-evoked postsynaptic currents, which was reversed by the addition of 4-aminopyridine (4-AP) (**Fig. 5g & h**). Together, these results show the existence of monosynaptic excitatory inputs from vSub to AHN^Vgat+^ neurons.

To examine whether AHN-projecting vSub neurons are responsible for AHN^Vgat+^ neuron activation during anxiogenic situations, we expressed GCaMP6s in vSub neurons projecting to AHN. This was achieved by injecting retroAAVs ^37^ encoding Cre-mCherry unilaterally into AHN of wildtype mice and AAVs encoding Cre-inducible GCaMP6s into vSub on the ipsilateral side (**Fig. 5i**). By monitoring GCaMP6s signals, we found that AHN- projecting vSub neurons displayed characteristic ramping activity during the approach-retreat bout in response to an unfamiliar object in the open field that aligned similarly to that found in AHN^Vgat+^ neurons (**Fig. 5j**). Furthermore, AHN-projecting vSub neurons showed higher and progressively increasing GCaMP6s signals in the open arm than the closed arm during the EPM test (**Fig. 5k-l**). Taken together, the close correspondence between the activity patterns of AHN- projecting vSub neurons and AHN^Vgat+^ neurons supports that vSub inputs drive AHN^Vgat+^ neurons in anxiety-provoking situations.

### Inhibiting AHN-projecting vSub neurons diminishes anxiety-related avoidance behavior

Finally, to test whether AHN-projecting vSub neurons acutely regulate anxiety-related avoidance behavior, we specifically inhibited these neurons by bilaterally injecting retroAAVs encoding Cre-mCherry into AHN and AAVs encoding Cre-inducible GtACR1 into vSub (**Fig. 6a-b**). Through *ex vivo* patch-clamp recordings, we confirmed that blue light pulses effectively and reversibly silenced GtACR1-expressing vSub neurons (**Fig. 6c-d**). By inhibiting the activity of AHN-projecting vSub neurons during the mouse’ approach towards the unfamiliar object in the open field (**Fig. 6e**), we completely abolished object-induced avoidance behavior in GtACR1 animals (**Fig. 6f-h**). Similarly, light inhibition of AHN-projecting vSub neurons also significantly increased the total time that the mouse spent in the light-illuminated open arm (**Fig. 6i-j**). This inhibition of open arm avoidance was also more evident in the second 5 min than the first 5 min of the trial (**Fig. 6k**). Together, these experiments establish that the activity of AHN-projecting vSub neurons is required for anxiety-related avoidance behavior.

**Fig. 6.**
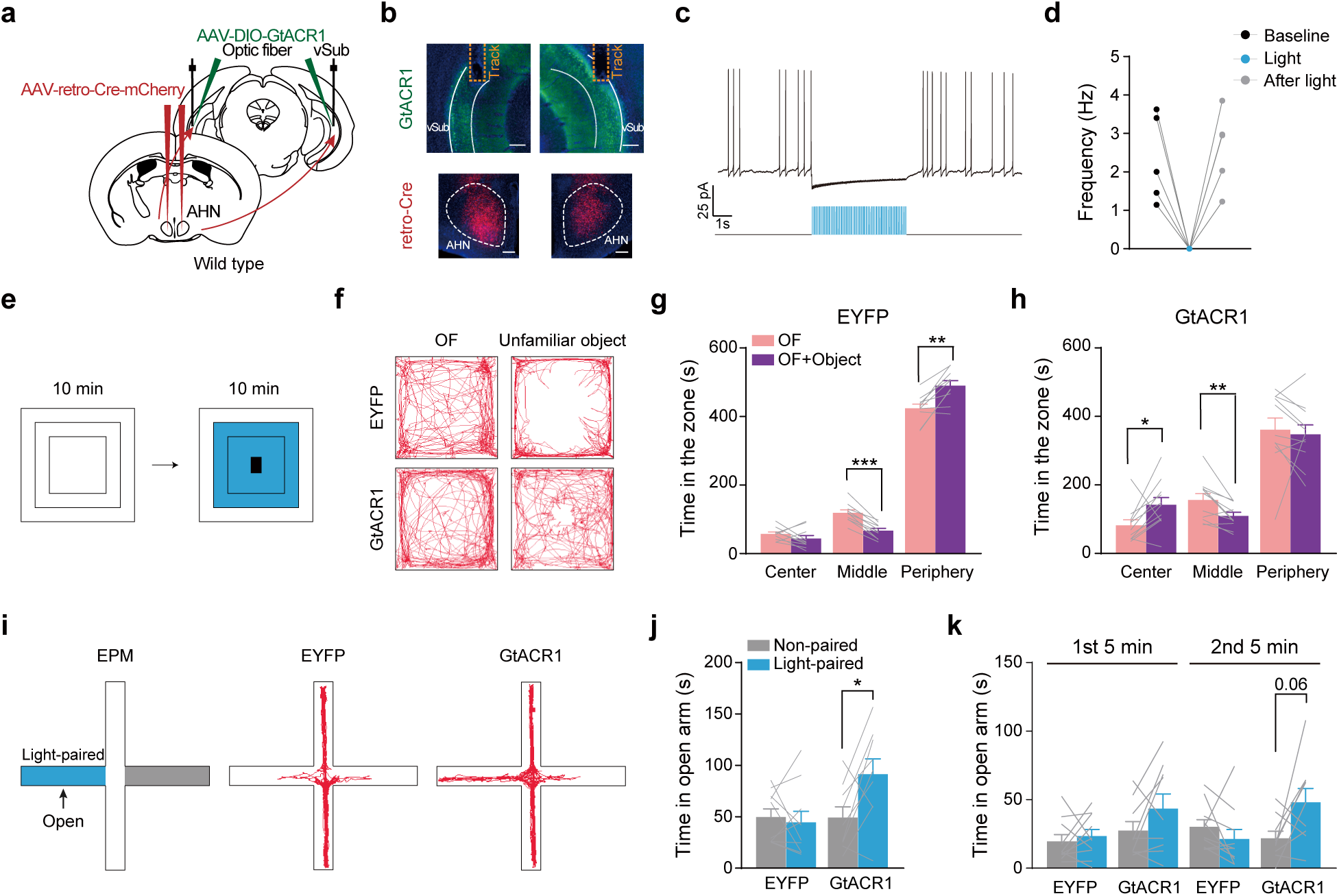
Inhibiting AHN-projecting vSub neurons diminishes anxiety-related avoidance behavior. (**a**) Schematics of the strategy to retrogradely target and bilaterally inhibit AHN-projecting vSub neurons. (**b**) Representative images showing GtACR1 expression in vSub (top), and retro-Cre expression in AHN (bottom). Scale bar, 200 μm. (**c**-**d**) A representative trace (c) and quantifications (d) showing trains of light pulses (473nm, 20ms, 20Hz), shown as blue lines in (c), acutely and reversibly inhibit firing of GtACR1-expressing cells. (**e-h**) GtACR1-mediated inhibition of AHN-projecting vSub neurons in open field. n = 10 EYFP and 11 GtACR1 males. (e) Schematics of light delivery restricted to the center and middle zone after object introduction. (f) Example open field trajectories of an EYFP and a GtACR1 male. (g-h) Time spent in the center, middle and periphery zones before and after object introduction with light stimulation. (**i-k**) GtACR1-mediated inhibition of AHN-projecting vSub neurons on EPM. n = 10 EYFP and 9 GtACR1 males. (i) Left, schematics showing light delivery restricted to one open-arm; right, example trajectories from an EYFP or a GtACR1 animal. (j-k) Time spent in the light-paired and non-paired open arm. *, p < 0.05; **, p < 0.01; ***, p < 0.001.

## Discussion

This study delineated a vSub to AHN pathway essential for anxiety-related avoidance behavior in two different conditions. As AHN is generally thought to be a node of the “predator defense circuit”, this vSub to AHN pathway provides a potential circuit mechanism to explain the well-known interaction between anxiety and predator defense behaviors. Moreover, the vSub-to-AHN circuit may channel mental assessment of potential threats by the hippocampus and other cognitive brain areas to initiate motor programs for avoidance. Such a pathway would allow for flexible, context-dependent, and individually varied displays of anxiety-related avoidance behaviors. Our results thus support the notion that anxiety is evolutionarily rooted in predator defense, as proposed by the “threat imminence” theory of anxiety behaviors ^18, 19, 21^.

Although AHN consists of predominantly *Vgat+* neurons, it is a heterogenous and ill-demarcated nucleus implicated in multiple behaviors such as thermoregulation, agonist behaviors, and predator defense ^38, 39^. Our results now present a new function for AHN^Vgat+^ neurons in mediating anxiety-related behavioral avoidance, but the exact identity of these neurons among the heterogenous AHN^Vgat+^ populations remains to be determined. In particular, we have defined an anxiogenic function for AHN^Vgat+^ neurons downstream of the vSub. However, some AHN inhibitory neurons receiving inhibitory projections from lateral septum Crfr2-expressing neurons may also inhibit stress-induced anxiety behaviors ^12^. Thus, there may co-exist anxiolytic and anxiogenic AHN^Vgat+^ neuronal populations representing subtypes of AHN neurons that serve opposite functions, analogous to the two subtypes of striatal medium spiny neurons expressing dopamine receptor 1 or 2 ^11, 40^. In addition, AHN local circuits may exist to link anxiolytic and anxiogenic neurons for modulating the approach vs. avoidance behavior in anxiety-provoking situations, similar to the local inhibitory microcircuits found in the central amygdala (CeA) for fear-related behaviors ^41, 42^. Thus, detailed characterization of AHN^Vgat+^ neurons in terms of transcriptional heterogeneity, activation patterns, and local- and long-range connectivity at single-cell resolution is of interest to further dissect the hypothalamic circuits for anxiety-related behaviors.

Psychologists have long postulated that the hippocampal formation is a center for computing, comparing, and arbitrating “safety” and “threat” signals to coordinate approach vs. avoidance in anxiety-provoking situations ^2, 43, 44^. Rodent studies have consistently shown that the ventral hippocampus, particularly vCA1, regulates anxiety-related behaviors in various paradigms ^9, 45–50^. The vCA1 neurons receive inputs from the amygdala ^51^ and project to separate several downstream targets, including the medial prefrontal cortex (mPFC), the lateral hypothalamus (LH), the lateral septum (LS), and BNST, to promote either approach or avoidance ^48, 50, 52^. Notably, the AHN-projecting vSub neurons we identified are located posteriorly and anatomically distinct from vCA1 neurons. As the output of the hippocampal formation, however, these vSub neurons are likely to receive direct inputs from vCA1 ^53^. Notably, we found that AHN-projecting vSub neurons showed progressively increasing activity on EPM, suggesting cumulation of internal regulatory signals during the behavior test. To our knowledge, such a progressive activation pattern has never been reported for anxiety-regulating neurons.

Our retrograde tracing study showed that AHN^Vgat+^ neurons receive inputs from mPFC, LS, LH, vSub, and BNST, all of which are projection targets of vCA1 neurons. Thus, AHN^Vgat+^ neurons may reside in a network position to integrate threat or safety-related signals transmitted and processed by these brain regions to initiate behavioral avoidance in anxiogenic situations. Importantly, we observed that AHN^Vgat+^ neuron activity tracks closely with avoidance behavior rather than the type of threats, and its level of increase showed individual specificity across different test conditions. Thus, AHN^Vgat+^ neurons may provide an entry point for understanding how excessive avoidance of perceived harm could emerge in some vulnerable individuals, such as psychiatric patients ^54^. In short, our results offer new insights into neural circuit mechanisms underlying anxiety-related behavioral avoidance.

## Methods

### Animals

All animals used in the study were adult males aged between 8-30 weeks. Wild-type males of C57BL/6J background were purchased from Shanghai SLAC Laboratory Animal Co., Ltd or Beijing Vital River Laboratory Animal Technology Co., Ltd. *Vgat-IRES-Cre* (*Slc32a1tm2^(cre)Lowl^/J*, Cat# 016962) was purchased from Jackson Laboratory. The animals were housed with *ad libitum* food and water under a reversed 12:12 hr light-dark cycle in the animal facility at the Institute of Neuroscience, excepted for those used in single-unit recording experiments, which were group-housed and bred in the animal facility at the Wuhan National Laboratory. Each cage contained at most six mice. Experiment protocols were approved by the Animal Care and Use Committee of the Institute of Neuroscience, Chinese Academy of Sciences, Shanghai, China (IACUC No. NA-01602016) or by the Hubei Provincial Animal Care and Use Committee and the Animal Experimentation Ethics Committee of Huazhong University of Science and Technology (IACUC No.844F).

### Virus

AAV-EF1α-DIO-mCherry (Serotype 2/8, titer 4.40 x 10^12^ vg/mL, vector genome per mL) and AAV-hSyn-ChR2-mCherry (Serotype 2/8, titer 8.40 x 10^12^ vg/mL) were purchased from Obio Technology Co, Shanghai. AAV-CAG-DIO-GtACR1 (Serotype 2/8, titer 2.20 x 10^12^ vg/mL) was purchased from Taitool Bioscience, Co, Shanghai. AAV-CAG-DIO-GtACR1 (Serotype 2/5, titer 5.00 x 10^12^ vg/mL) was purchased from PackGene Biotech Co, Guangzhou. AAV-EF1α-DIO-H2B-EGFP (Serotype 2/8, titer 8.33 x 10^12^ vg/mL), AAV-EF1α-DIO-EYFP (Serotype 2/8, titer 3.58 x 10^12^ vg/mL), AAV-hSyn-DIO-GCaMP6s (Serotype 2/8, titer 4.80 x 10^13^ vg/mL) and AAV-retro-hSyn-cre-mCherry (Serotype 2/2, titer 7.00 x 10^13^ vg/mL) were purchased from gene editing core facility of Institute of Neuroscience. AAV-EF1α-DIO-RVG (Serotype 2/9, titer 2.00 x 10^12^ vg/mL), AAV-EF1α-DIO-EGFP-2A-TVA (Serotype 2/9, titer 2.00 x 10^12^ vg/mL) and RV-EnVA-DG-DsRed (2.00 x 10^8^ IFU/mL, infectious units per mL) were purchased from BrainVTA, Wuhan.

### Mouse surgery

Surgeries were performed as previously described ^55^. Stereotaxic surgeries were performed on a David Kopf Model 1900 frame or a custom-built frame (Cat# SH-01, Xinglin LifeTech) that allows brain targeting at an angle. The animals were anesthetized with 0.8 - 5% isoflurane or with intraperitoneal (i.p.) injection of 1% pentobarbital sodium and hypodermic injection of 5 mg/kg carprofen for pain relief. The coordinates used for viral injection were based on the Paxinos and Franklin Mouse Brain Atlas, 2nd edition. For unilateral targeting of the AHN, coordinates of AP: - 0.820 mm, ML: ± 0.500 mm, DV: -5.200 mm were used. For bilateral targeting of the AHN related to optogenetic inhibition experiments, the coordinates were adjusted to be AP: - 0.820 mm, ML: ± 1.400 mm, DV: -5.100 mm at an angle of 10 degrees. For targeting the vSub, the coordinates were AP: - 4.100 mm, ML: ± 3.650 mm, DV: -3.800 mm. ∼ 60 - 200 nl of the virus was injected into the target brain site with a home-made nano-liter injector (Cat# SMO-10, Xinglin LifeTech) at a flow rate of ∼ 70 nl/min. Optic fibers (diameter, 200 mm; N.A., 0.37; Hangzhou Newdoon Technology Co.,Ltd) were implanted ∼50 μm above the viral injection site and secured onto the skull for fiber photometry recordings with dental cement and skull screws. For optogenetic inhibition, optic fiber was implanted 300 - 500 μm above the injection site. Animals were allowed to recover at least three weeks before being tested in behavioral experiments. For pseudorabies tracing experiment, ∼ 80 - 150 nl of the 1:1 mixture of helper virus (AAV-DIO-TVA-GFP and AAV-DIO-RG), or ∼ 100 nl AAV-DIO-TVA-GFP alone for control experiments, was first injected unilaterally in the AHN of *Vgat-IRES-Cre* mice and three weeks later, ∼ 100 - 150 nl RV-EnVA-DG-DsRed into the exact location. Histological analysis was carried out 1 week later. Ovariectomized (OVX) surgeries were performed with animals anesthetized with i.p. injections of ketamine (80 mg/Kg) and xylazine (8 mg/Kg), and animals were allowed to recover for over one week after the surgery prior to subsequent experiments.

### Histology

Histological analysis was performed as previously described ^56, 57^. Briefly, animals were anesthetized with 10% chloral hydrate and perfused with PBS, or DEPC treated PBS followed by 4% PFA. Brains were post-fixed overnight in 4% PFA at 4 °C and sectioned at 40 μm using a vibratome (VT1000S, Leica) except for experiments involving the RNAscope kit (ACD Bio.). All virally expressed fluorescent proteins or fusion proteins were visible without immune-staining. All brain sections were counterstained with DAPI (Sigma, Cat# d9542, 5mg/ml, 1:1,000). Images were captured by a 10 X objective fluorescent microscope (Olympus, VS120) or confocal microscope (Nikon, C2). For pseudorabies virus tracing, brain sections were evenly divided into two sets and only one set was mounted, imaged with a 10 X microscope (Olympus, VS120), and processed in ImageJ software. dsRed+ cells were counted outside of the AHN injection site and assigned to specific brain areas according to the Allen Institute adult mouse coronal atlas (http://atlas.brain-map.org/). The percentage inputs (% inputs) was calculated for each injection site by dividing the number of dsRed+ cells found in each brain region by the total number of dsRed+ cells tallied.

For histological analysis involving the RNAscope kit, after perfusion and post-fix, brains were dehydrated with 30% sucrose in DPEC-PBS and sectioned at 20 μm using a microtome and mounted onto SuperFrost Plus® Slides (Fisher Scientific, Cat. No. 12-550-15). RNA probes for *Vgat* (Cat #319191) and *Vglut2* (Cat #319171-C3) were ordered from ACD Bio. The *in situ hybridization* was performed using the RNAscope kit (ACD Bio.), following the user manual. Two brain sections covering the AHN were selected from each mouse. Images were captured with a 20X objective using a confocal microscope (Nikon C2) and processed in ImageJ software. Based on the DAPI counter-staining signal, the numbers of *Vgat* and *Vglut2* neurons in the AHN were counted.

For validating the *Vgat-IRES-Cre* line, we used RNAscope Fluorescent Multiplex Assay combined with immune-fluorescent staining. One brain section was selected from each mouse. After the *in situ*, brain slices were blocked by 2.5% BSA (Sigma Cat #V900933) for an hour, then stained overnight at 4°C with chicken anti-GFP antibody (ABCAM, Cat #ab13970, dilution 1:300). The next day, the brain sections were rinsed three times with 1 X PBS before incubating with the secondary antibody, goat-anti-chicken Alexa 488 (Jackson Immuno Research Laboratories, Cat #103-545-155, dilution 1:300) for two hours. Images were captured with 60X objective using a confocal microscope (Olympus FV3000) and processed in ImageJ software. Three 400 x 400-pixel squares were selected from each brain section, analyzed, and quantified for the proportion of co-labeled neurons.

### Behavioral tests

Mice were singly housed two days before behavioral experiments and were handled once per day for these two days. Animals were continuously singly housed during the period of behavioral tests. All behavior tests were recorded with a camera at a frame rate of 25 or 30 Hz. For behavioral tests in the home cage, a stimulus, an object or a hormonally primed ovariectomized female, was introduced after the animal was moved to the video-taping area to acclimate for ∼ 10 mins. For the open field (OF) test, mice were introduced into a corner of 40 x 40 x 40 cm white box under illumination, ∼ 10 min after which, either an object (unfamiliar or familiar) was introduced into the OF or the experimenter’s hand was put briefly about the box mimicking the motion of object introduction. Afterward, behaviors were recorded for another 10 min. The unfamiliar object used included type C battery, acrylic cuboid cube, toy airplane and metal paper clip, presented on separate testing days in a pseudo-randomized manner. The familiar object used was a type C battery co-housed for three days with the tested animal. For behavioral analysis, the OF box was divided into three zones. The “center” zone encompasses the innermost 20 x 20 cm square; the “peripheral” zone is the region within 5 cm along the wall, and the rest the “middle” zone.

For single unit recording, a mouse was introduced to an open field arena and allowed to first explore for ∼ 5 min. Afterward, an unfamiliar object was introduced into the center and the mouse was monitored for another 5 - 10 min. Next, the mouse was introduced to a clean cage (30 x 20 x 20 cm) and allowed to explore for 5 min. Then a semicircular filter paper (7 cm diameter) spotted with ∼ 400uL red fox urine (Lenonlures company, USA) was introduced to one side of the cage and the mouse was monitored for another 5 - 10 min. The EPM (Elevated Plus-Maze) apparatus used consists of a central region (5 x 5 cm), two open-arms (30 x 5 cm), and two close-arms (30 x 5 x 15 cm), in a “+” configuration and placed 50 cm above the floor. At the beginning of the EPM test, the mice were put in the center area oriented towards a close-arm.

All behavioral videos were annotated with custom-written MATLAB code as previously described ^56^. Approach start was defined as the mice headed to and began to move toward the object and the end as the mice retreating from the object, which was also the start of the retreat. Retreat end was scored when animals stopped moving. A social investigation was defined as nose-to-face and nose-to-body contacts initiated by the male towards the female, sniff was defined as nose-to-urogenital contact, and mount was defined as male placing its forelimbs on the back of the female and climbing on top. The time that animals spent in each OF zone or EPM arm were extracted with EthoVision XT (Noldus) or custom-written MATLAB code. Example trajectories were generated in EthoVision XT (Noldus).

### Fiber photometry

Fiber photometry recordings were carried out as previously described ^56^. Before the recording, the implanted optic fiber was connected to the recording device (Biolink Optics Technology Inc., Beijing) through an external optic fiber. Briefly, 488 nm laser was reflected through a dichroic mirror (MD498, Thorlabs), and the fluorescence signal was passed through a bandpass filter (MF525-39, Thorlabs) and collected in a photomultiplier tube (PMT, R3896, Hamamatsu). Emission signals were low-pass filtered at 30 Hz and sampled at 500 Hz with a data acquisition card (USB6009, National Instrument) using software provided by Biolink Optics. A LED bulb was transiently triggered at the start of the recording session to facilitate alignment of the fiber photometry recording signal and animal behaviors for data analysis.

For data analysis, fluorescent signals acquired were analyzed with custom-written MATLAB code. Briefly, raw signals were first adjusted according to the overall trend to account for photo-bleaching. Afterward, the values of fluorescence signal change (ΔF/F) were calculated as (F−F_0_)/F_0_. In this formula, F represents the signal value at any given moment, and F_0_ represents the baseline. For recordings done in the open field, F_0_ was the average signal value over the 10 mins after the animals were placed in the open field and before the object introduction. When tested in the home cage, F_0_ was the average fluorescence value over 10 mins before introducing stimulus (an object or a female). For the EPM test, F_0_ was the mean fluorescence value for the 10 mins recording period. To calculate the ΔF/F value for a defined open field zone or EPM location, we first extracted the body location of the mice in each frame to assign the ΔF/F value to a specific zone or location and then averaged ΔF/F values of all frames that belonged to a particular zone. To align ΔF/F signals with behavior, ΔF/F values were segmented based on behavior events and averaged first across different events in a trial and then across different trials from each animal.

To calculate the correlation between the GCaMP6s signal and approach-retreat bout in Figure 1G, we first excluded the behaviors with an inter-behavioral interval of less than one second. We then transferred the trend-adjusted F value and the behavior data into a binary (0 or 1) form and calculated the correlation between the two using a nonparametric Spearman correlation test. For GCaMP6s signal, we defined time points with an F value over two standard deviations (2 SD) away from the mean as ‘‘1’’ and otherwise as ‘‘0’’. For approach-retreat behavior, any time points annotated with the behavior was ‘‘1’’ and otherwise as ‘‘0’’. For all correlation analyses involving approach end ΔF/F signal (Figure 1J-1N), we excluded recordings of the first approach from the analysis to rule out any possible effects of initial exposure. We used a nonparametric Spearman correlation test to calculate the correlation between the approach end ΔF/F signal and the approaching interval in Figure 1J, 1M. Other correlation analysis of ΔF/F signal and behavioral time were calculated using the parametric Pearson correlation test. Heatmap representations of ΔF/F value on EPM were generated with custom-written MATLAB code.

### Single-unit recording

Single-unit recording was performed and analyzed as previously ^58, 59^. Briefly, the guide tubes housed 16-channel electrodes of 25.4-mm formvar-insulated nichrome wire (Cat # 761500, A-M System, USA). The final impedance of the electrodes was 700–800kU. Mice were implanted with the 16-channel electrodes targeting AHN and then allowed to recover for at least five days before further behavioral tests. Before the testing, mice were singly housed and connected to the recording connector for two days to adapt. During the recording, the 16-channel electrodes were connected to an amplifier and sampled by a computer. Recorded signals were amplified (3 200 000 gain) and digitized at 40 kHz by the NeuroPhys Acquisition System (Neurosys 2.8.0.8, USA) and NeuroLego System (Jiangsu Brain Medical Technology Co.ltd). Raw signals were filtered (300 - 6000 Hz) to remove field potential signals. Single-unit spike sorting was performed using the MATLAB toolbox (MClust-4.4). Waveforms with amplitudes smaller than 50 - 60 uV (three times noise band) were excluded from the analysis. Unsorted waveforms were analyzed with peak value and two types of principal components. We manually defined waveforms with similar characters into clusters. A cluster of waveforms was considered a single neuron if the ratio of its inter-spike-interval (ISI) under 2ms was less than 1%, the isolation distance was greater than 20, and L-ratio less than 0.1 ^60, 61^. In addition, if the spike time of any two units coincided via the cross-correlation comparison, those units were also considered a single unit.

The neuron firing rate was analyzed by first extracting the spike train frequency during the 5 s before and after an approach towards an unfamiliar object or a sniffing bout of fox urine. Data was binned by 250 ms. Object-approach and fox urine-sniff responses were calculated as Z-scores by normalizing the 20 bins of during-behavior firing rates to the 20 bins of before-behavior baseline firing rates. Neurons with a Z-score >2 (p < 0.05) during any two consecutive bins within the 2 s after the onset of the behavior were classified as excited neurons, whereas neurons with a Z-score <-2 (p < 0.05) were classified as inhibited neurons.

### Optogenetic inhibition

Before the test, the bilateral optic fibers were connected to a 473 nm laser power source (Shanghai Laser and Optics Century Co. or Changchun New Industries Optoelectronics Tech Co., Ltd.). Light delivery was controlled by LabState (AniLab), which detects the centroid of the animal in real-time to trigger the laser or turn it off. In the OF test, the light was triggered when the centroid of the animal entered the center and middle zone immediately after object introduction or when the centroid of the animal entered the center zone starting 10 mins after object introduction, for 10 mins. For the second scenario, animals were monitored for another 10min after cessation of light stimulation as the “post-light” stage. For the EPM test, the light was triggered when the centroid of the animal entered one randomly selected open-arm immediately after the animal was placed on the EPM or when the centroid of the animal entered either of the two open-arms 10 mins after the animal was placed on the EPM, for a duration of 10 mins. For the second scenario, animals were monitored for another 10mins after cessation of light stimulation as the “post-light” stage. For real-time place preference, the apparatus used consists of two 17 x 17 cm chambers and a 5-cm-wide gap in between the two chambers. One chamber was black with a metal-rod floor, and the other chamber was white with a wire floor. The light was triggered whenever the centroid of an animal entered a randomly chosen light-paired chamber. Light power in all these experiments was 5 mW, 20 Hz, 20 ms.

### Brain slice electrophysiological recording

Mice were anesthetized with isoflurane, perfused transcardially with ice-cold oxygenated (95% O_2_/5% CO_2_) high-sucrose solution (in mM, 2.5 KCl, 1.25 NaH_2_PO_4_, 2 Na_2_HPO_4_, 2 MgSO_4_, 213 sucrose, 26 NaHCO_3_). Brains were sectioned coronally at 250 μm using a vibratome (Leica, VT1200S) in an an ice-cold oxygenated high-sucrose solution. Brain sections containing the AHN or vSub were incubated in artificial cerebrospinal fluid (in mM, 126 NaCl, 2.5 KCl, 1.25 NaH_2_PO_4_, 1.25 Na_2_HPO_4_, 2 MgSO_4_, 10 Glucose, 26 NaHCO_3_, 2 CaCl_2_) at 34 °C for 1 hr. The intracellular solution for recordings contains (in mM) 135 K-gluconate, 4 KCl, 10 HEPES, 10 sodium phosphocreatine, 4 Mg-ATP, 0.3 Na_3_-GTP and 0.5 biocytin (pH:7.2, 276 mOsm). Recording electrodes (3-5 MΩ, Borosilicate Glass, Sutter Instrument) were prepared by a micropipette puller (Sutter Instrument, model P97). For synaptic transmission recordings, repetitive single pulses of blue light (10 ms, power 12 mW/mm^2^) were delivered onto the brain slice through a 40×objective with an X-Cite LED light source (Lumen Dynamics). Cells were clamped at 0 mV for IPSC recording and at -70 mV for EPSC recording. To validate the mono-synaptic connections between vSub and AHN neurons, 1 μM of tetrodotoxin (TTX, absin, Cat# abs44200985a) and 1 mM 4-aminopyridine (4-AP, Alomone Labs, Cat# A-115) were sequentially added into the bath solution. To confirm the effects of neuronal inhibition by GtACR1, repetitive 20 Hz pulses of blue light ( 20 ms, power 7 mW/mm^2^, interval 20 s) were delivered onto the AHN or vSub brain slice. Whole-cell recordings were performed using a MultiClamp700B amplifier and Digi-data 1440A interface (Molecular Devices). Data were low-pass filtered at 2 kHz and sampled at 20 kHz under voltage clamp, while low-pass filtered at 10 kHz and sampled at 10 kHz under current clamp. All experiments were performed at 33 °C with a temperature controller (Warner, TC324B).

### Statistical Analysis

Statistical tests were analyzed with GraphPad Prism (GraphPad Software). For comparisons between two groups, we first analyzed the distribution of the data with the Shapiro-Wilk normality test. Data sets that passed the normality test were analyzed with Student’s t-test (two-tailed, paired, or unpaired); otherwise, we used the Wilcoxon matched-pairs signed-rank test for paired data and used non-parametric Mann–Whitney U-test for unpaired data. For comparisons among data of more than two groups, such as in Fig. 2C, one-way ANOVA was used. All data were plotted as mean ± standard error of the mean (SEM). *, p < 0.05; **, p < 0.01; ***, p < 0.001.

## Acknowledgements

We thank the Drs. Mu-ming Poo, Xiao-Ke Chen, Ning-Long Xu, Zhe Zhang, Ji Hu for their valuable comments on the manuscript. This work was supported by grants from the Strategic Priority Research Program of the Chinese Academy of Sciences (XDB32000000) and the National Nature Science Foundation of China (31871066, 31922028, 31900721), and by Shanghai Municipal Science and Technology Major Project (Grant No. 2018SHZDZX05).

## Author Contributions

J.J.Y. & X.H.X. designed the experiments, analyzed the data and wrote the manuscript. A.X.C. analyzed RV tracing result, prepared the figures and scored behavioral videos. W.Z. performed electrophysiological recordings. T.H. performed single-unit recordings with the help from M.H. under the supervision of H.L. X.J.D. performed behavioral tests. Y.Z.X. performed *post-hoc* immunostaining of brain slices. Y.L. Zhang maintained the mouse colonies.

## Data and material availability

All data and material are available upon publication and personal request.

## Extended Data Figure Legend

**Extended Data Fig. 1.**
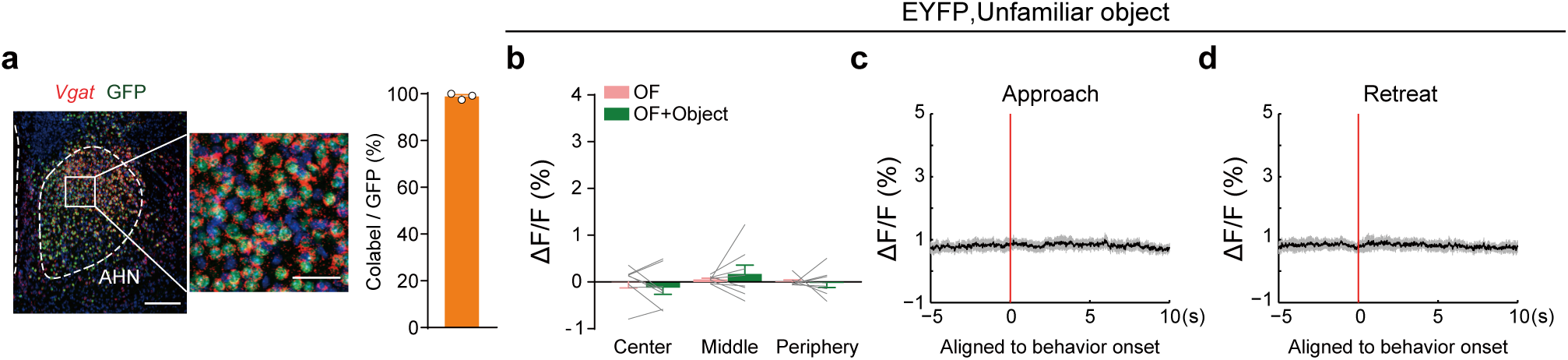
Verification of the *Vgat-IRES-Cre* mouse line and fiber photometry recordings in control EYFP male mice. (**a**) AAV-EF1α-DIO-H2B-EGFP was injected into AHN of *Vgat-IRES-Cre* males. A representative image on the left shows *Vgat in situ* hybridization signals and viral-mediated GFP expression in AHN. Scale bar, 200 μm. The magnified image on the right highlights the area within the white box. Scale bar, 50 μm. Quantification shows the co-localization of *Vgat* and GFP signals. n = 3 mice. (**b-d**) Fiber photometry recordings of EYFP mice in open field with an unfamiliar object. n = 8 mice.

**Extended Data Fig. 2.**
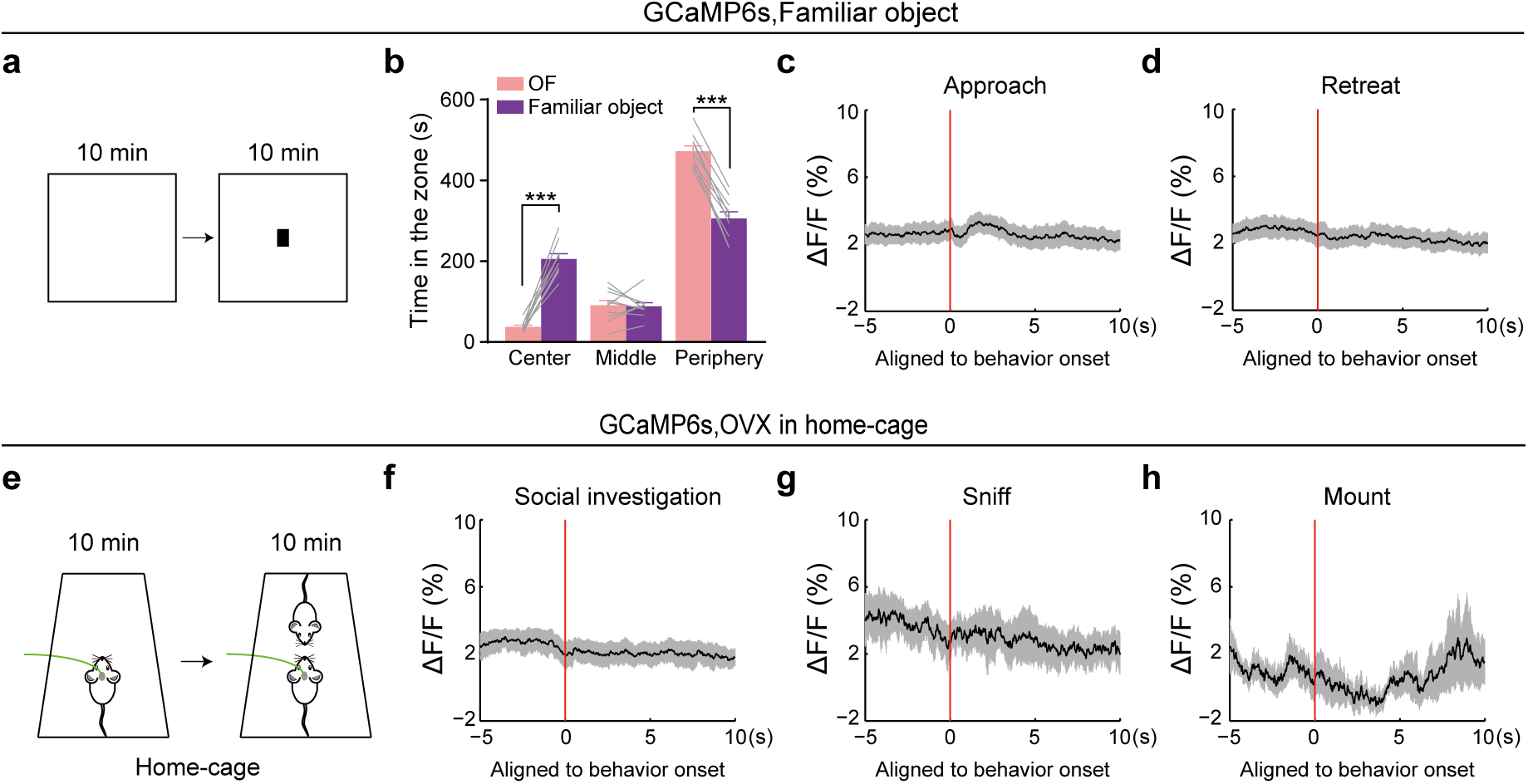
Fiber photometry recordings of AHN^Vgat+^ activity in response to a familiarized object in an open field and to a female conspecific in the homecage. (**a-d**). Fiber photometry recordings of GCaMP6s males with a familiarized object. n = 10 mice. (a) The object (a battery) used was placed in the mouse’s homecage for three days before introduced to the open field. (b) Quantification of the time the mice spent in each zone of the open field before or after introduction of the familiar object. Mice spent significant time in the center zone after object introduction. (c-d) Average values of GCaMP6s ΔF/F signal aligned to approach (c) or retreat onset (d) at the time “0”. Shades indicate the SEM. No changes in AHN^Vgat+^ activity was detected during either behavior. (**e-h**) Fiber photometry recordings of GCaMP6s males interacting with an unfamiliar, hormonally primed ovariectomized (OVX) female mouse in the home cage. n = 9 mice. (e) Schematics of the behavioral protocol. No changes in AHN^Vgat+^ activity was detected during social investigation (f), sniff (g), or mount (h). ***, p < 0.001.

**Extended Data Fig. 3.**
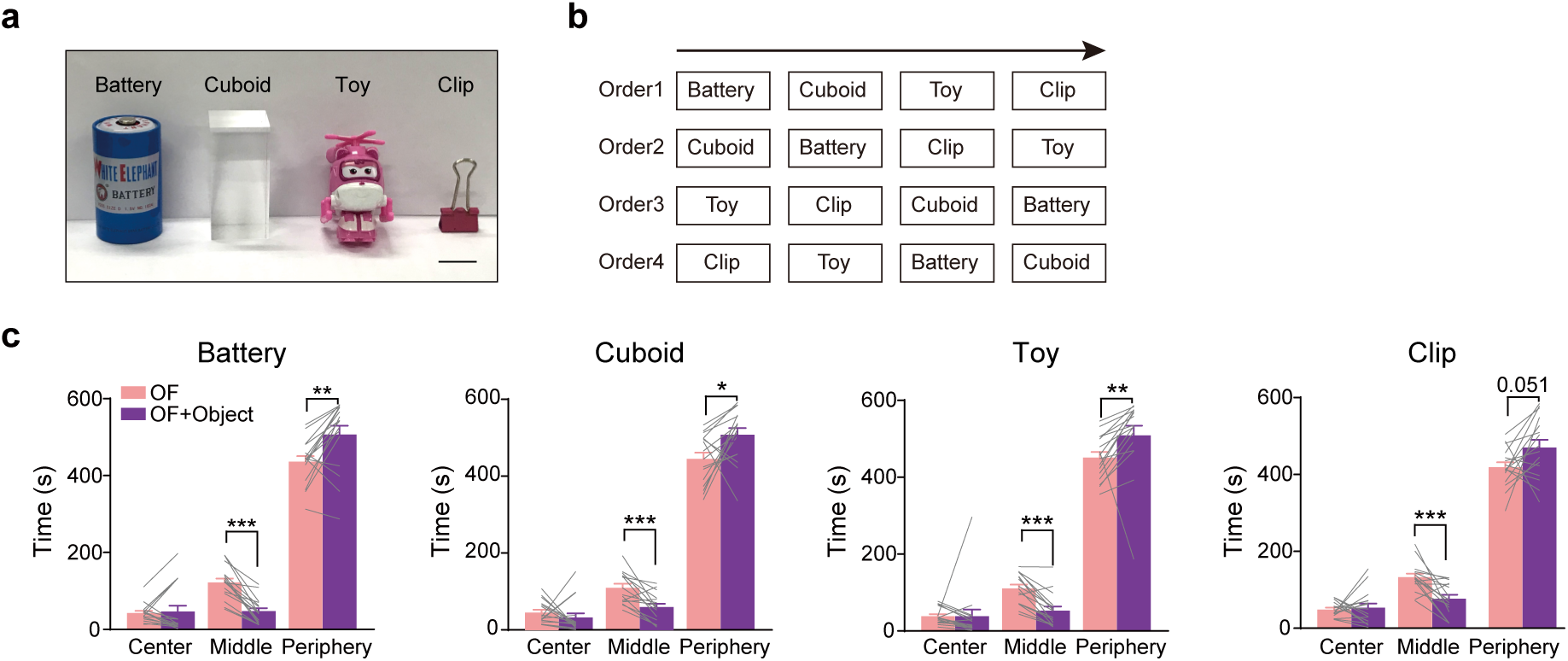
Different objects induced similar center avoidance and periphery preference in the open field test. **(a)** Different unfamiliar objects used. **(b)** The order in which the unfamiliar objects were presented on separate testing days. **(c)**Time spent in the center, middle, and periphery zone of the open field before or after the indicated object was introduced. n = 16 mice. *, p < 0.05; **, p < 0.01; ***, p < 0.001.

**Extended Data Fig. 4.**
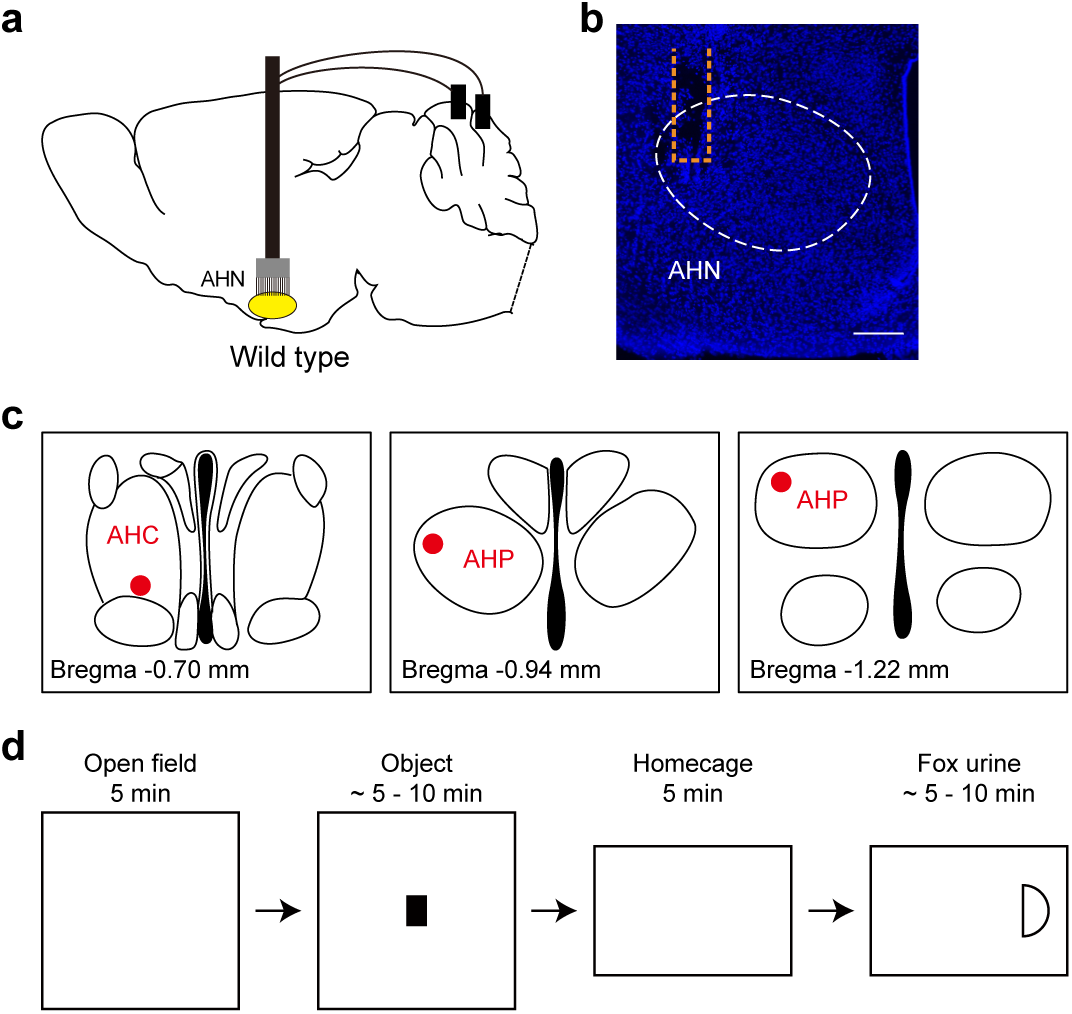
Single-unit recordings of AHN neurons. **(a)** Schematics showing electrode implantation in AHN and grounding of implanted electrodes. **(b)** A representative *post-hoc* image showing the tip of the implanted electrode lied within AHN. Scale bar, 200 μm. **(c)** Anatomical tip locations of the implanted electrode in the three recorded mice. **(d)** Behavioral procedures of single-unit recording experiments.

**Extended Data Fig. 5.**
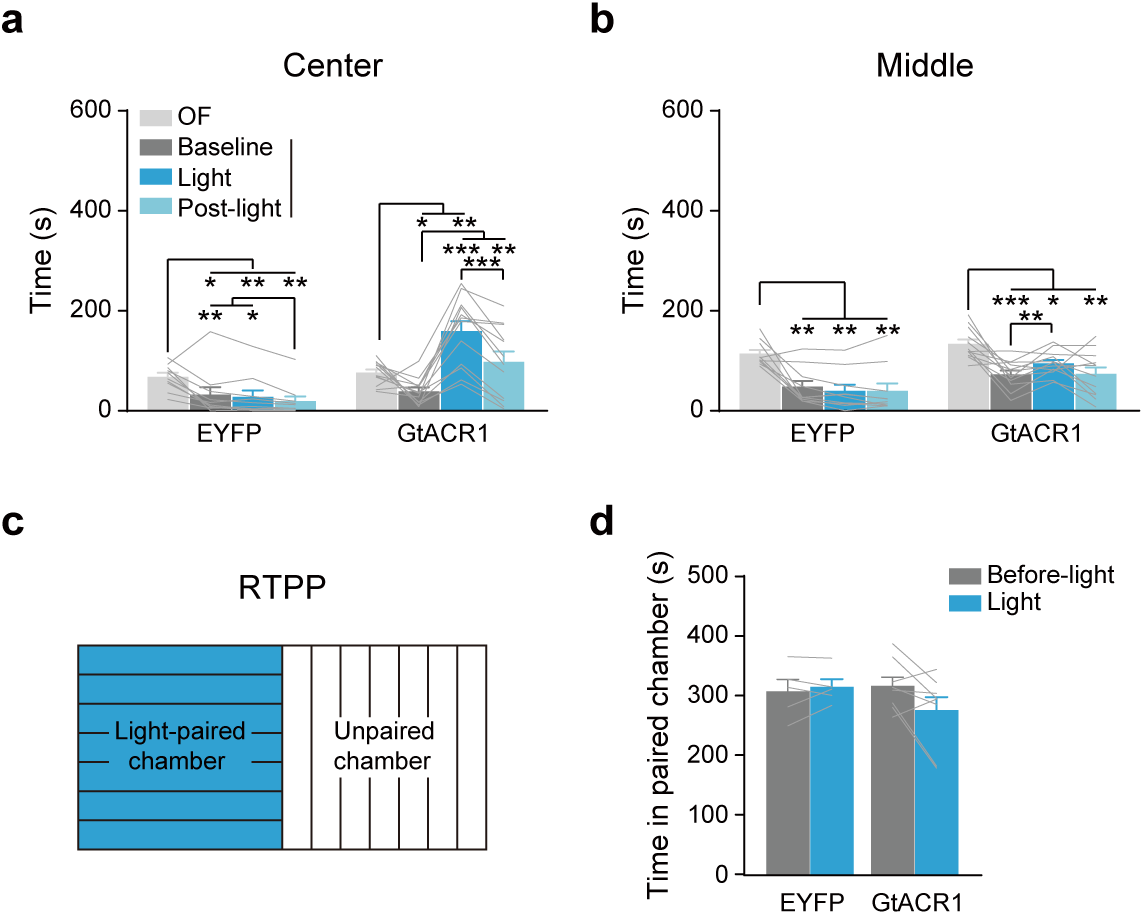
Optogenetic inhibition of AHN^Vgat+^ neurons reduces object-induced center avoidance in open field but does not lead to conditioned place preference. (**a-b**) Time spent in the center zone (a) and middle zone (b) in open field test before or after an object introduction. n = 10 EYFP and 12 GtACR1 males. (**c**) Schematics of the real-time place preference test. The blue region indicates the light-paired chamber and the other unpaired chamber. (**d**) Time spent in the light-paired chamber before or during light stimulation, 10 min each. n = 5 EYFP and 8 GtACR1. *, p < 0.05; **, p < 0.01; ***, p < 0.001.

**Extended Data Fig. 6.**
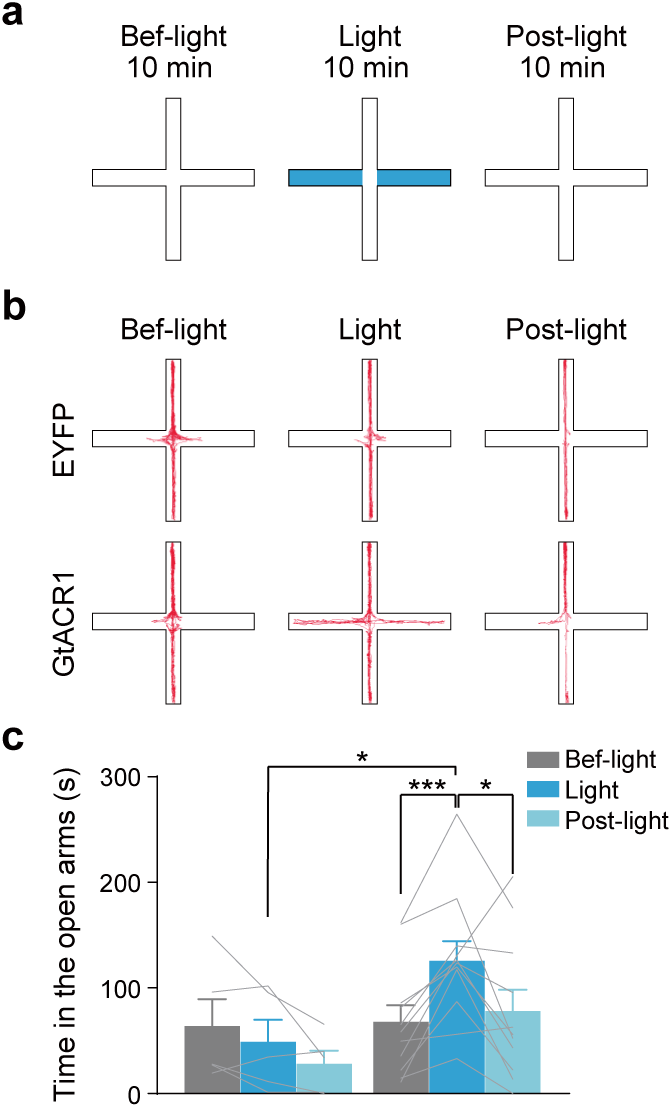
Optogenetic inhibition of AHN^Vgat+^ neurons reduces EPM open arm avoidance. (**a**) Schematics of the light delivery patterns and timing. **(b)** Example movement trajectories on EPM from a control EYFP and a GtACR1 male. (**c**) Time spent in EPM open arm in before, during, and post-light delivery. Light illumination increased open arm time in GtACR1 but not control EYFP males. n = 5 EYFP and 11 GtACR1 males. *, p < 0.05; ***, p < 0.001.

**Extended Data Fig. 7.**
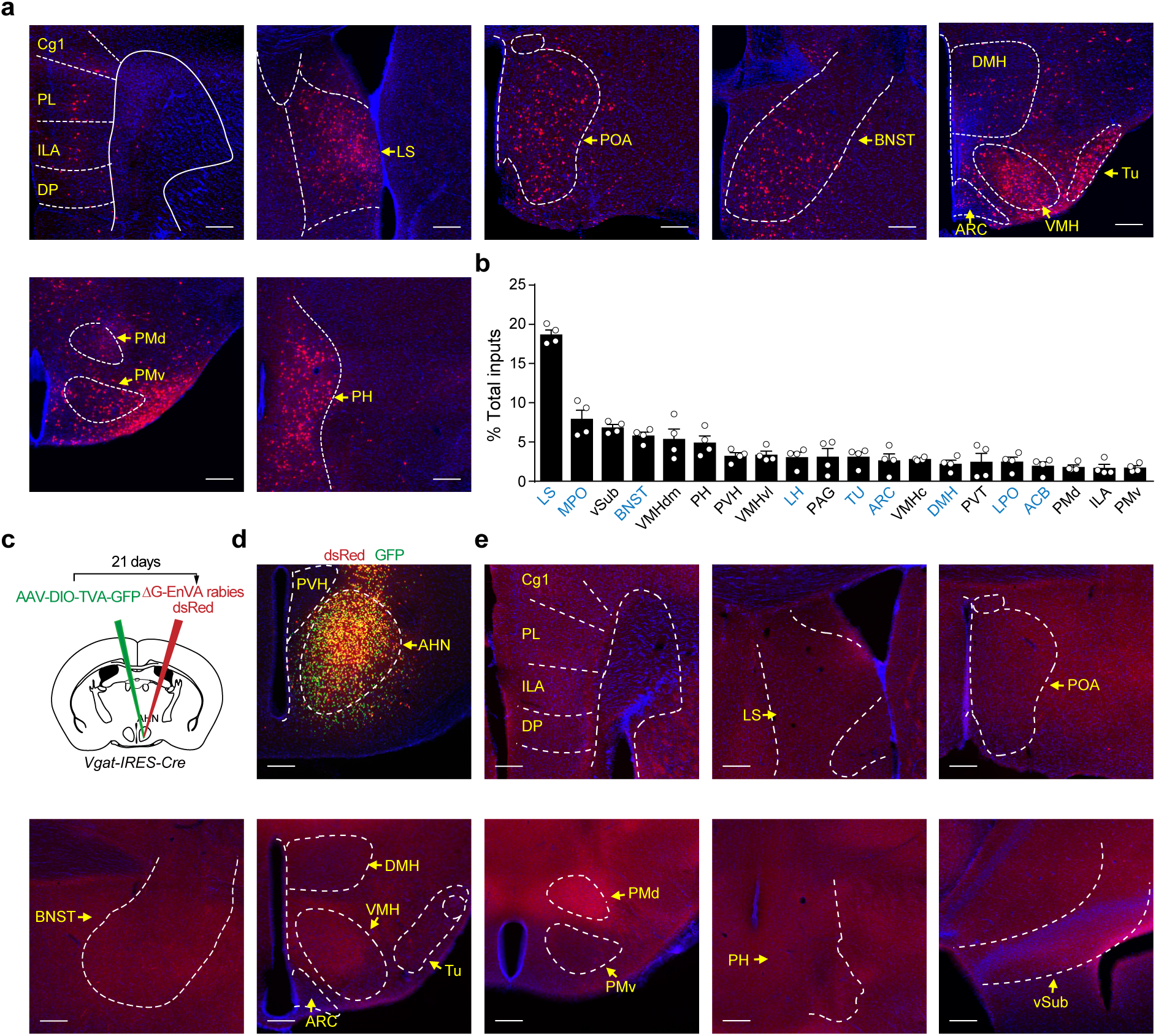
Quantification of and control experiments for pseudorabies mediated retrograde tracing of inputs to AHN^Vgat+^ neurons. (**a-b**) Pseudotyped rabies virus-mediated retrograde tracing of inputs to AHN^Vgat+^ neurons. (a) Representative images showing dsRed+ neurons in areas indicated. Scale bar, 200 μm. (b) Quantification of dsRed+ neurons in each region as % of total dsRed+ cells detected outside of the AHN. n = 4 mice. Light blue text indicates areas consisting of predominantly inhibitory projection neurons (www.mouse.brain-map.org). (**c-e**) The control experiment. n = 3. (c) Schematics of the viral strategy for the control experiment without RG injection. (d) A representative image showing infection of AHN^Vgat+^ neurons by AAV-DIO-TVA-GFP and EnVA-pseudotyped rabies virus expressing dsRed. Scale bar, 200 μm. (e) Representative images showing no dsRed+ signal in areas indicated. Scale bar, 200 μm. Abbreviations: cingulate cortex area 1 (Cg1), prelimbic area (PL), infralimbic area (ILA), dorsal peduncular area (DP), lateral septum (LS), preoptic area (POA), paraventricular hypothalamic nucleus (PVH), bed nuclei of the stria terminalis (BNST), dorsomedial hypothalamus (DMH), ventromedial hypothalamus (VMH), arcuate hypothalamic nucleus (ARC), tuberal nucleus (TU), dorsal premammillary nucleus (PMd), ventral premammillary nucleus (PMv), posterior hypothalamus (PH), ventral subiculum (vSub).

